# Resources modulate developmental shifts but not infection tolerance upon coinfection in an insect system

**DOI:** 10.1101/2024.08.01.606236

**Authors:** Nora K.E. Schulz, Danial Asgari, Siqin Liu, Stephanie S.L. Birnbaum, Alissa M. Williams, Arun Prakash, Ann T. Tate

**Affiliations:** Department of Biological Sciences, Vanderbilt University, Nashville TN 37232; Evolutionary Studies Initiative, Vanderbilt University, Nashville TN 37232

## Abstract

Energetic resources fuel immune responses and parasite growth within organisms, but it is unclear whether energy allocation is sufficient to explain changes in infection outcomes under the threat of multiple parasites. We manipulated diet in flour beetles (*Tribolium confusum*) infected with two natural parasites to investigate the role of resources in shifting metabolic and immune responses after single and co-infection. Our results suggest that gregarine parasites alter the within-host energetic environment, and by extension juvenile development time, in a diet- dependent manner. Gregarines do not affect host resistance to acute bacterial infection but do stimulate the expression of an alternative set of immune genes and promote damage to the gut, ultimately contributing to reduced survival regardless of diet. Thus, energy allocation is not sufficient to explain the immunological contribution to coinfection outcomes, emphasizing the importance of mechanistic insight for predicting the impact of coinfection across levels of biological organization.

## Introduction

If we know how individual stressors like infection affect traits and dynamics at a given biological scale, can we predict how multiple stressors will function in tandem? To unite and explain broad trends of species abundance, persistence, and ecosystem functioning in the face of stress and change, ecological theory leans heavily on metabolic and stoichiometric models that rely on assumptions about the flow and use of resources and energy at different scales of biological organization (Ott *et al*. 2014; Bernot & Poulin 2018). For example, resource-focused theory has promoted recent advances in our understanding of ecology within organisms and the maintenance of microbiomes, symbionts, and parasites (Rynkiewicz *et al*. 2015). Whether drawing on simplified resource allocation assumptions or more complex dynamic energy budgets, within-host models have generated testable predictions for infection outcomes and the oscillation and persistence of parasites (Cressler *et al*. 2014; Ramesh & Hall 2023) as they directly or indirectly compete with immune systems for resources. As these frameworks grow in popularity, however, it is worth asking about the extent to which resource allocation can fully explain infection outcomes, particularly in the face of multiple stressors such as coinfection.

Hosts in nature are likely to endure exposure to parasites at multiple periods in their lives; coinfection occurs when these exposure events lead to infection with two (or more) parasite species simultaneously (Tate 2019). Since both species exploit host resources and generally induce or modulate immune responses, they can facilitate or antagonize each other and lead to infection and transmission outcomes that differ from single infection scenarios (reviewed in (Rovenolt & Tate 2022)). To what extent can resource theory explain these outcomes? This question largely depends on the relative sensitivity of parasites, immune dynamics, and damage repair to resource conditions within the host (Graham 2008; Clay *et al*. 2023). After all, resource limitation can alter allocation among life history traits and reconfigure immune system investment (Adamo *et al*. 2016), altering the reception of incoming parasites and their transmission potential (Vale *et al*. 2013). Sometimes, resource availability dominates parasite competition from the bottom-up regardless of immunological regulation. In mice, for example, gut nematodes destroy red blood cells and thereby limit malaria parasite propagation through resource limitation even though the nematodes also suppress immune responses, which should otherwise facilitate parasite replication and transmission (Griffiths *et al*. 2015). On the other hand, helminth co-infection in African buffalo stimulates a T cell polarization state away from the optimal regime needed to fight coccidia. While it is not clear – and indeed not probable – that this immune shift is primarily resource-driven, it does lead to a more hospitable gut habitat for the coccidia that ultimately increases parasite shedding at the (presumably energetic) expense of host reproduction (Seguel *et al*. 2023).

The resource sensitivity of these molecular mechanisms can be more directly tested in experiments that manipulate resource availability. Recent examples of coinfection outcomes in insect systems demonstrate limited (Deschodt & Cory 2022) or mixed effects (Zilio & Koella 2020) of resource limitation on host and parasite fitness-associated traits, but these studies focus on infection outcomes rather than the mechanisms that drive them. To what extent does a primary infection alter the metabolic and immunological landscape encountered by a second parasite species? Does resource allocation ultimately drive these differences, or should we be selective about generalizing coinfection dynamics from energy budget models?

To test these questions, we turned to a model system for host-parasite population biology – the confused flour beetle *Tribolium confusum* (Park 1948) and two of its natural parasites. The first parasite, the eugregarine *Gregarina confusa*, induces a chronic but avirulent infection that provides tractability for these questions by avoiding confounding effects of morbidity and mortality (Detwiler & Janovy 2008; Thomas & Rudolf 2010). The second parasite is the entomopathogenic bacterium *Bacillus thuringiensis* (Bt), which instigates acute mortality (Behrens *et al*. 2014) and is sensitive to the immune dynamics of the host (Jent *et al*. 2019). We divided larvae in a factorial design that included presence or absence of gregarine exposure and a standard or nutrient-limited diet. To quantify the extent to which the primary parasite influences the metabolic and immune landscape experienced by the second parasite, we used mRNA-seq to investigate molecular signatures of metabolic and immunological shifts to gregarine infection and measured diet-dependent development time and metabolite levels in gregarine-infected or uninfected larvae. We then infected larvae with Bt and investigated transcriptomic signatures of gregarine infection on immune and metabolic trajectories during acute Bt infection. We applied these results to disentangle the relative impact of resources and immunity on gut pathology, infection resistance, and disease-induced mortality. Our results suggest that while some coinfection-induced shifts in life history parameters may be approximated with energy budget assumptions alone, others are largely insensitive to resources and rely instead on shifts in pathology associated with immune responses and damage repair. Since these parameters are particularly important for predicting population dynamics and parasite-mediated apparent competition outcomes at the community level (Johnson *et al*. 2015; Cortez & Duffy 2020; Rovenolt & Tate 2022), ecologists should account for basic immunology before relying too heavily on resource theory for coinfection models.

## Materials and Methods

### Beetle rearing, handling, and diet

*Tribolium confusum* beetles for this experiment were derived from a stock colony collected in 2013 from Pennsylvania, USA (Tate & Graham 2015) and subsequently kept under laboratory conditions (standard diet, 30C, in the dark). To create breeding groups for the experiments, we took 60-80 adults per group from a colony and allowed them to lay eggs in 16g flour for 24 hours. We then combined eggs for all breeding groups and distributed them across experimental diets. Besides age-matched eggs from the same egg-laying period, we also created staggered breeding groups two days apart to derive larvae of different ages but equivalent sizes.

The *standard diet* for 200 larvae consists of 16g autoclaved whole-wheat flour (Fisher) and 5% w/w brewer’s yeast (Fleischmann) in a 100mm petri dish. To reduce the protein and nutrient quality of the diet we excluded the yeast (*no-yeast diet*). Yeast is an important source of protein and other nutrients, and larvae raised on this restricted diet will still develop into adulthood but generally more slowly. In these experiments, all diets were further modified by reducing new flour to 10g and adding 6g flour derived from gregarine-infected or clean mini-colonies as described below.

### Infection protocols

To create infectious flour, we added beetles from gregarine-infected or uninfected stocks to clean flour for four days, providing time to deposit infectious gregarine oocysts. We then removed the beetles and used the flour to create the diets (gregarine exposure: 4g infected flour + 2g uninfected flour; no exposure: 6g uninfected flour). This method resulted in a 60%+ prevalence of gregarine infection in all experiments, as determined by gut microscopy (Thomas & Rudolf 2010), (q)PCR (f: CCTCGAGGAAGTTCGAGTCTAT, r: TTGACAGCTTGGGCACTTTAT, 400nM efficiency = 99.2%, Tm = 55C), and/or deposited gametocyst counts 18-24 hours after temporary starvation (Janovy *et al*. 2007). Once ingested, gregarine trophozoites attach to the guts for 7-10 days before finding a mate, forming a gametocyst, and evacuating the gut (Janovy *et al*. 2007).

To challenge larvae with Bt, we produced bacterial cultures as described in (Jent *et al*. 2019). Plating of the infection culture confirmed a concentration of 1.8 *10^9^ CFU/mL, which results in an LD50 dose in *T. confusum*. To septically inoculate larvae, we dipped a micro-dissection needle in the bacterial (or saline control) aliquot and stabbed it into the space between the head and second segments.

### Development and metabolite assays and standardizing age vs stage + stats

We counted freshly emerged pupae and dead larvae daily from day 22-30 post oviposition and removed them to avoid resource-deprived larvae receiving additional protein from cannibalism (Park *et al*. 1970). To collect size-equivalent larvae for the metabolite measurements, we adjusted the collection dates according to the developmental delays in the different treatments, *i.e.* greg+/yeast+ and greg-/yeast- larvae were collected three days later than the greg-/yeast+ control treatment, while the slowest greg+/yeast- larvae were collected a week later.

To measure the primary metabolites, we froze size-matched larvae after starvation, washed them twice in cold insect saline, and homogenized them (n=18-21 larvae/diet-pathogen treatment). A third of each sample went into each of the three performed measurements. We measured total protein content in a Bradford assay (Schulz *et al*. 2023). For the glucose assay we used the GO Assay Kit (Sigma) and for lipids a Vanillin assay (Abcam) (Barr *et al*. 2023).

We analyzed differences in larval development time to pupation among the diet and gregarine treatment groups (N = 77-150 larvae/treatment) using log-rank survival analyses from the “survival” package in R (R_Core_Team 2012; Therneau 2014). We used linear models to confirm treatment-wise mass equivalence among larvae selected for metabolite assays (Bates D. 2010). After standardizing lipid, glucose, and protein measurements by individual larval mass, we used generalized linear models with gamma distributions (due to positive values and some right-skew) to evaluate differences among parasite and diet treatments and their interactions for each metabolite (N = 17-64 larvae per treatment).

### Survival assays

To analyze Bt infection-induced mortality of gregarine-infected and uninfected larvae under different resource conditions, we raised larvae from age-staggered breeding groups on contaminated standard or no-yeast diets, size-standardized larvae from each treatment (because smaller larvae generally have higher mortality regardless of treatment), and then infected the beetles with an LD50 dose of bacteria (N = 48 Bt-infected and 8 saline-challenged larvae/treatment/block for 4 experimental blocks). Bt induces rapid-onset mortality, generally between 8 and 14 hours post infection (Jent *et al*. 2019). We excluded larvae that died early from the trauma of inoculation, and then monitored mortality from 6-14 hours, performing a final check at 24 hours as most larvae will have died or recovered from Bt infection by then (Tate *et al*. 2017). We analyzed larval survival using Cox proportional hazards (coxme package in R (Therneau & Therneau 2015)).Proportional hazards assumptions were not met (tested using the coxzph function) because survival rates among Bt-infected and saline control beetles were so drastically different. Therefore, we stratified by infection treatment;gregarine exposure, diet, and their interaction served as main effects and experimental block as a random effect.

### Gut integrity assays

To determine whether the damage or immune responses instigated by gregarine parasites accelerate mortality, we evaluated the gut barrier integrity of beetle larvae. We randomly assigned eggs from breeding groups to the four gregarine-by-diet treatment groups in 96-well microplates. After 20 days, we septically exposed larvae to an LD20 dose (1.6 x10^6^ CFU/mL) of Bt or mock-infected them with insect saline. Subsequently, we placed the larvae on blue dye food prepared with 2.5% FD&C blue dye no.1 (Spectrum Chemicals) using a protocol described by (Zanchi *et al*. 2020). After 20 hours of feeding, we examined the distribution of blue food dye under a microscope to detect any leakage in the gut-intestine barrier, scoring the beetles exhibiting a blue “smurf” phenotype. We analyzed smurf proportions using binomial generalized linear models with block as a main effect and then gregarine status, diet, and Bt infection status as main and interacting effects.

### Time series and RT-qPCR analysis of parasite loads

To evaluate the impact of gregarine infection on host-Bt dynamics, we first collected gut samples (N = 6 pools of 8 guts/gregarine treatment) from gregarine-exposed or clean larvae. We then challenged larvae from these groups with a needle dipped in sterile saline (control) or the experimental dose of live Bt. Beetles were sacrificed every two hours for the 12 hours of the acute infection phase (n = 8-12 Bt-infected and 6 uninfected larvae/time point) and stored individually at -80C. We extracted RNA using Qiagen RNeasy mini-kits, confirmed RNA concentration using the Nanodrop, and then reverse-transcribed RNA into cDNA (VILO mastermix). We quantified Bt load via RT-qPCR (SybrGreen) as previously described (Jent *et al*. 2019; Critchlow *et al*. 2024). We validated our qPCR primers (see above) on known gregarine infected and uninfected samples to devise a threshold of detection and used these primers to categorize gregarine-exposed samples as currently infected or not. We log-transformed the linearized dCt values for normality (Jent *et al*. 2019) and used linear models in R (“lm” function) to analyze the impact of Bt exposure, time, gregarine exposure or confirmed infection, and the interaction of time and gregarines on relative Bt loads. Because bacterial loads bifurcate over time, and variation in high-load beetles might not be captured across the entire load distribution, we also used a Bt load threshold on samples from 6-12 hours post infection to characterize beetles as high-load, and performed logistic regression (lme4 package, glm function, family = binomial and link = logit) on high-load status vs gregarine exposure/infection. The results were not sensitive to the chosen threshold or on whether larvae were merely exposed to gregarines or actively infected.

### Transcriptome assembly and annotation

In addition to the gut samples, we chose whole-body larval RNA samples from the time series (sample sizes and time points in **Table S1**) that exhibited a near-median bacterial load for the treatment and time point, to avoid introducing load-induced variance (Tate & Graham 2017). 150bp paired end libraries were produced using the Illumina TruSeq kit and sequenced in a single batch at the Vanderbilt VANTAGE core on the Illumina NovaSeq 6000 (complete statistics in **Table S2**). Sequencing data is publicly available on NCBI Sequence Read Archive (accession PRJNA771764). We first assessed RNAseq read quality using fastqc (Andrews 2010). There is currently no published annotated genome for *T. confusum* so using only samples not infected with gregarines, we assembled a *de novo* transcriptome using Trinity with default settings (contig statistics in **Table S3**); quality filtering was performed within Trinity and reads were assembled in paired-end mode (Grabherr *et al*. 2011). Highly similar transcripts were clustered using cd-hit (Fu *et al*. 2012). To assess the quality of the assembly, the reads were realigned to the assembled transcriptome using bowtie2 and the ExN50 statistic was calculated within Trinity (Grabherr *et al*. 2011). To assess the completeness of the assembly, transcripts were analyzed using BUSCO (Benchmarking Universal Single-Copy Orthologs, **Table S4**) against an insect gene set (Manni *et al*. 2021).

We used kallisto v 0.48.0 to quantify gene expression (Bray *et al*. 2016) by performing pseudo alignment of RNA-seq reads to the assembled transcriptome of *T. confusum* and summing count or transcript per million (TPM) values across isoforms. Because we were interested in achieving a high degree of accuracy for AMP-specific analyses, we also used Coleoptera AMPs as training sets (**Table S5**) and constructed Hidden Markov Model (HMM) profiles using HMMER (Finn *et al*. 2011) to annotate AMPs in the *T. confusum* proteome. Since some of our analyses relied on annotation data from a well-developed genome (Herndon *et al*. 2020), we filtered bit scores to combine results from BlastP and Blastx to identify *T. confusum* orthologs of *T. castaneum* genes (**Figs. S1**, **S2**). Full methodological details for analytical pipelines and sample processing are described in the Supplementary Methods.

### Differential expression analyses

We used the DEseq2 (v 1.36) package in R (Love *et al*. 2014) to run three differential expression (DE) analyses. In the first analysis, we identified DE genes in the gut upon gregarine infection relative to uninfected guts (sample details in **Tables S1,2**). In the second and third analyses, we identified DE genes upon Bt or coinfection in the whole body samples at six (N=4) or eight (N = 6) hours post infection with Bt relative to uninfected beetles (N = 8) or beetles infected only with gregarines (N=8). Within each time point, we modeled differential gene expression in DEseq2 as expression ∼ Bt status + gregarine status + their interaction (false-discovery rate (FDR) corrected *P-*value < 0.05 (Benjamini & Hochberg 1995)). Because we were concerned about type II error after FDR adjustment due to the large number of annotated but low-expressed *T. confusum* genes, we also used the seSeq (2.30) package in R (Hardcastle & Kelly 2010) to investigate the AMPs specifically. To this end, we divided samples into four groups: genes that are not DE across samples, genes that are DE in samples infected by Bt, genes that are DE in the gregarine-infected samples, and genes that are DE in co-infected samples relative to other samples, and reported the posterior probability of differential expression. We tried identifying TF binding sites upstream of relevant AMP genes but we have low confidence in the accuracy so the results are not presented here.

We performed weighted gene co-expression network analysis (WGCNA (Langfelder & Horvath 2008)) on genes that have orthologs in *T. castaneum* to identify modules of co-expressed genes. Genes with zero count values across all replicates were removed before analyses. Next, we constructed a signed correlation matrix for each analysis (merging threshold = 0.25, minimum module size = 30) using the count data and used them to identify positive or negative correlations between the expression of genes in the network. We calculated the Pearson correlation of the module eigengenes across samples to identify modules of co-expressed genes. Using the associated *T. castaneum* ortholog gene ids, we performed gene ontology (GO) analyses with DAVID (Huang *et al*. 2007) to find functional categories for co-expressed modules identified via WGCNA. In addition, we performed Kyoto Encyclopedia of Genes and Genomes (KEGG) analyses on differentially expressed genes using clusterProfiler package in R (Yu *et al*. 2012).

## Results

### Both restricted diet and gregarine infection prolong development time, but hosts can compensate metabolically unless stressed by both

In isolation, both yeast restriction (Fig. 1A, Log-rank test, N=150, developmental hazard ratio (HR) = 0.76(0.6-0.96), p = 0.024) and gregarine infection (N=82, HR = 0.73(0.55-0.97), p = 0.029) significantly and equivalently prolonged larval development time by approximately one day (Fig. 1B) relative to well-fed and uninfected reference larvae (N = 128). This effect was exacerbated when gregarine infection and yeast restriction were combined, leading to significantly slower development times than all other treatments (N = 77, HR relative to reference = 0.46(0.35-0.62), p < 0.001).

**Figure 1.**
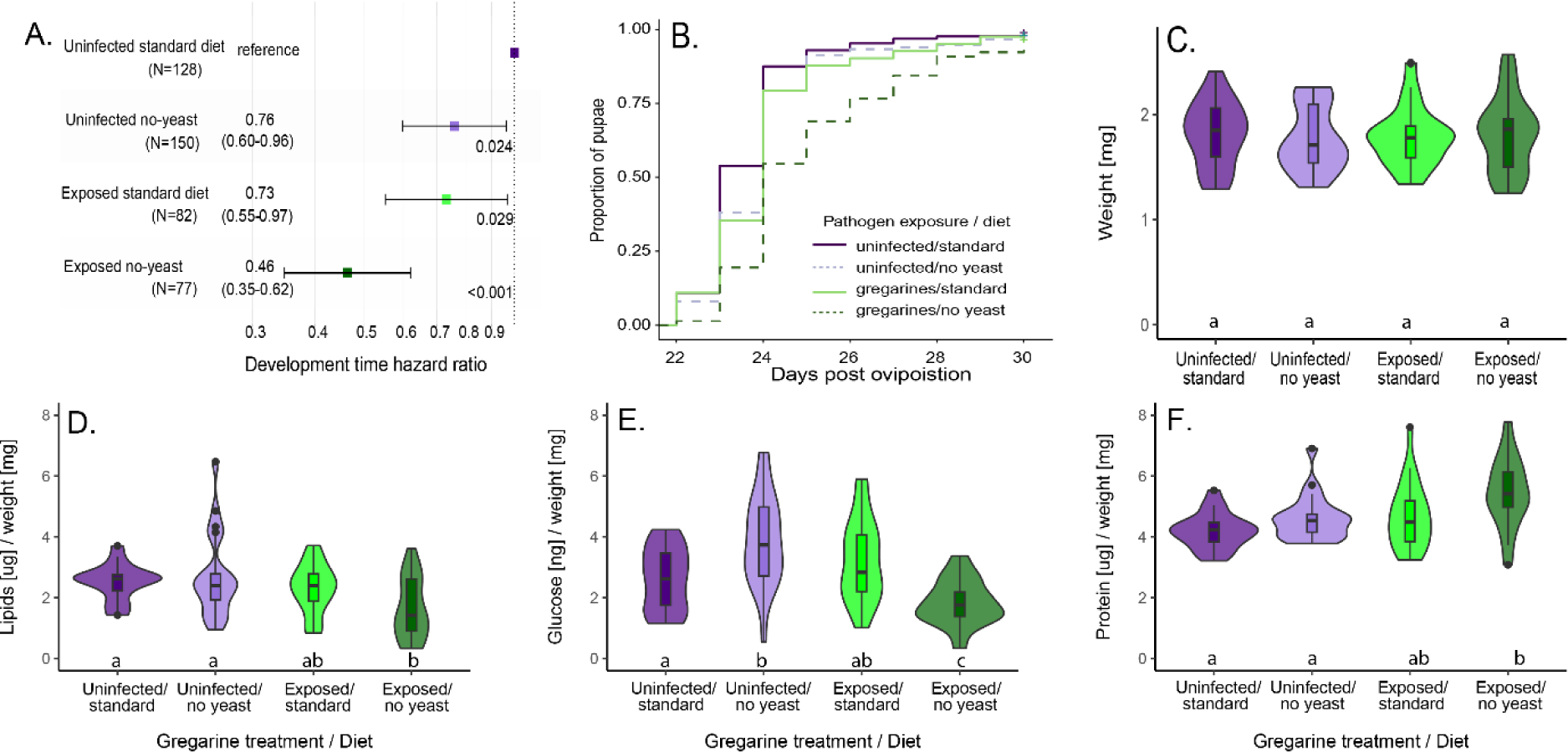
Effects of chronic gregarine infection, resource limitation, and their interaction on larval development and metabolic profiles. A) Log-rank statistics (hazard ratio of development time to pupation, 95% CI, and p values; note that a smaller HR means they develop more slowly) and B) development curves for the rate of larval development to the pupal stage for gregarine-infected larvae under standard or yeast-restricted diets relative to uninfected larvae on standard diets. In mass-standardized larvae (C), the effect of gregarine infection and diet on lipid (D), glucose (E), and protein content (F) were analyzed using GLMs with gamma distributions (Table 1); post-hoc pairwise test (BH-corrected) bins appear in lowercase letters.

**Table 1.** The impact of diet, gregarine infection, and their interaction on within-host metabolites in mass-matched larvae.

| *Mass* | estimate | se | t | p |
| --- | --- | --- | --- | --- |
| No yeast/uninfected | 0.047 | 0.097 | -0.49 | 0.63 |
| Standard/infected | -0.03 | 0.097 | -0.31 | 0.76 |
| No yeast/infected | -0.011 | 0.097 | -0.11 | 0.91 |
| ***Lipid*** |  |  |  |  |
| Intercept | 0.926 | 0.100 | 9.3 | **<0.0001** |
| Gregarine exposure | -0.088 | 0.140 | -0.6 | 0.53 |
| No-yeast diet | 0.067 | 0.140 | 0.5 | 0.63 |
| Gregarine: No-yeast | -0.412 | 0.197 | -2.1 | **0.039** |
| ***Glucose*** |  |  |  |  |
| Intercept | 0.965 | 0.087 | 11.2 | **<0.0001** |
| Gregarine exposure | 0.216 | 0.122 | 1.8 | 0.082 |
| No-yeast diet | 0.392 | 0.122 | 3.2 | **0.0020** |
| Gregarine: No-yeast | -0.960 | 0.173 | -5.5 | **<0.0001** |
| ***Protein*** |  |  |  |  |
| Intercept | 1.423 | 0.051 | 28.0 | **<0.0001** |
| Gregarine exposure | 0.134 | 0.039 | 3.4 | **0.00064** |
| No-yeast diet | 0.121 | 0.039 | 3.1 | **0.0021** |

Notes: individual larval mass was analyzed using an lm evaluating all groups relative to the standard diet uninfected reference group. All metabolites were modeled using gamma-distributed glms and diet, gregarines, and their interactions as main effects. The interaction effect was dropped from the protein analysis as it was not significant and its exclusion yielded a better model fit.

To analyze metabolic profiles across the treatments, we minimized the confounding effect of development time discrepancies by controlling for larval mass (Fig. 1C; N = 21 per treatment; mass range 1.3-2.6mg; p > 0.6 among all pairwise treatment comparisons) rather than age or instar, which is indeterminate in flour beetles. Neither diet nor gregarine infection significantly affected mass-corrected lipid levels (Table 1), but the interaction of the two was significant, as the infected and no-yeast group had reduced lipid stores (Fig. 1D; interaction p = 0.027). Diet and its interaction with gregarines significantly predicted glucose levels in opposite directions (Fig. 1E), as no-yeast diet larvae had a significantly higher glucose level than the reference group (posthoc BH-corrected p = 0.003) but the infected and no-yeast group had significantly lower glucose levels (interaction p <0.0001). Protein (Fig. 1F) made up a significantly higher proportion of larval body mass in no-yeast (p = 0.005) and gregarine-infected groups (p = 0.002), likely indicating that a higher proportion of total mass is structural rather than stored resources; the interaction effect was not significant. These values were not significantly dependent on individual larval mass within treatments except for lipids in gregarine-exposed individuals (**Fig. S3**), which increased for larger larvae in the gregarine-infected standard-diet treatment and decreased in the gregarine-infected-low protein diet.

### Gregarine-infected guts reveal altered metabolic and immunological profiles

A principal component analysis (PCA) of transcriptomic profiles revealed a clear separation of gregarine-infected and uninfected gut samples (**Fig. S4**). Upon DESeq2 analysis, upregulated genes were enriched for ribosomal, cuticular, and glycolytic proteins, while downregulated genes included a bacterial recognition protein (GNBP-1) as well as metabolic and digestive enzymes such as apolipoproteins, lipases, trehalose transporters, cytochrome P450s, cathepsin B, α-L-fucosidase, carboxypeptidase A, and juvenile hormone binding proteins (**Table S6**). Zooming in on immune effector responses, we identified the significant differential regulation of four upregulated and tightly co-expressed AMPs in the gut upon gregarine infection ((Fig. 2A, **Table S7**) Defensin-1, Attacin-2, Attacin-3, and Cecropin-3) as well as two downregulated AMPs (Cecropin-1 and PR5-3, which contains a thaumatin domain; Fig. 2B).

**Figure 2.**
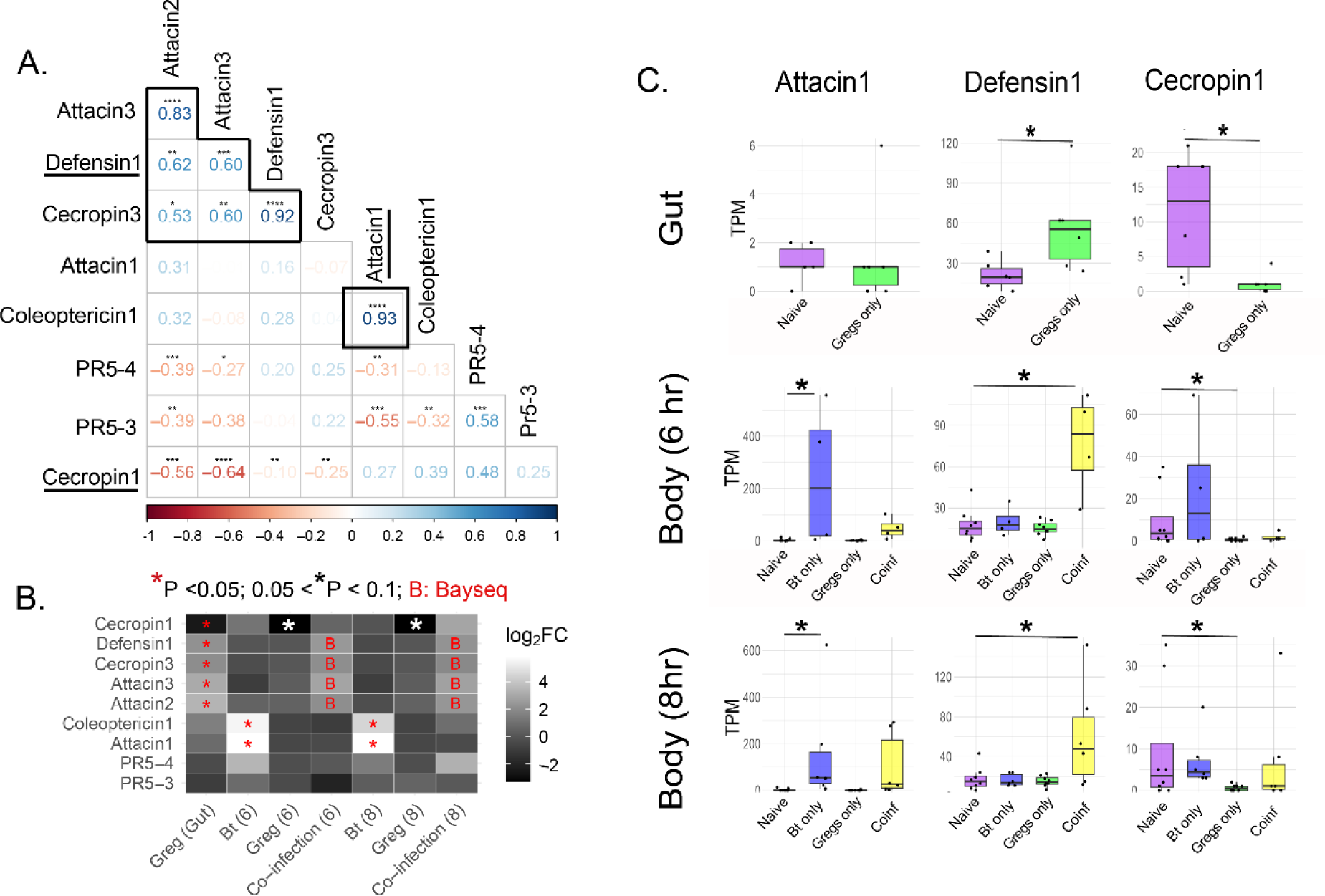
Effector protein (AMP) expression patterns upon infection with Bt, gregarines, and co-infection. A) The Pearson correlation matrix of expressed AMPs across all samples includes highlighted clusters of co-expressed effectors. B) Differential expression analysis of effectors across treatments showing log2-fold change of effectors in the gut and at 6 and 8 hours post Bt infection relative to naïve reference; DE genes delineated with asterisks. Four AMPs identified as DE upon co-infection using Bayseq are shown with red “B”. C) Expression patterns over time and treatments for three AMPs that each represent a cluster of co-expressed effectors (underlined in A). The Y-axis shows normalized counts (transcripts per million; TPM) and the x-axis shows treatments. Significant differences relative to naïve are shown with asterisks.

WGCNA analysis did not identify any specific gene co-expression modules that were significantly associated with gregarine infection (**Table S8**), but KEGG analysis revealed significant enrichment of the ribosome followed by non-significant hits on steroid synthesis and signatures of altered carbohydrate metabolism (**Table S9**).

### Transcriptomic profiles from gregarine-infected and coinfected larvae reveal altered physiology and unique immune responses to single vs coinfection

We evaluated transcriptomic profiles of whole-body samples from individual gregarine-infected or uninfected larvae six or eight hours post co-infection with *Bacillus thuringiensis* (**Table S1**). PCA showed effects of Bt or gregarine infection at six hours post-infection, but this separation became muddled by eight hours post-infection (**Fig. S4**).

Focusing first on immune-specific analyses, we discovered that two AMPs (Coleoptericin-1 and Attacin-1) were highly associated with the main effect of Bt infection at both six and eight hours post infection (Fig. 2C). Meanwhile, gregarine infection was associated with the downregulation of the same cecropin (Cecropin-1) originally downregulated in the gut, although this effect was muted in the coinfected samples (Fig. 2C). The same AMPs that were upregulated in the gut upon gregarine infection were also uniquely upregulated in the whole-body coinfection samples at both 6 and 8 hour time points (Fig. 1C; Defensin-1, Attacin-2, Attacin-3, and Cecropin-3).

While p*-*values for these four AMPs did not survive multiple corrections (adj *P* > 0.05) (**Tables S10, S11**, **Fig. S5**), we hypothesized that this is due to a type-II error on the FDR because of the unusually large number (>200k) of draft-annotated *T. confusum* genes. Therefore, we ran DE analyses using the Bayseq package and calculated the posterior probabilities instead of *P-*values (Fig. 2B), which dispenses with the need for multiple corrections. Posterior probabilities of upregulation due to co-infection for these genes were close to one (**Fig. S5**) but were larger at six hours post-infection compared to eight hours post-infection, suggesting that their expression is reduced after six hours (e.g. TPM count in **Fig. 2C**).

We ran co-expression network analyses to identify gene modules that are uniquely co-regulated according to each infection treatment (**Tables S12, S13**). We limited our analyses to genes with orthologs in *T. castaneum* because running analyses using the complete dataset is computationally intensive and because GO terms are not annotated for *T. confusum*. Our co-expression results at six hours post infection were generally consistent with our AMP-specific analyses, such that DE AMPs within each analysis belonged to the same modules (e.g., *Attacin-1* and *Coleoptericin-1* both belong to the purple 6 hour WGCNA module; **Table S14**), and the expression of genes in each module was consistent with the direction of differential expression identified by DEseq2 (**Table S10**). Two modules showed significant associations with Bt (purple) and co-infection (cyan) (**Fig. S6**), and the expression of genes within these modules was almost exclusively linked to these specific treatments. The top GO and individual DE terms associated with Bt infection included cell adhesion proteins and wound healing/immune defense (e.g. hemocyanin activity, mucins, PGRP-SC2, AMPs, serine proteases and serpins; **Table S15**). KEGG analyses indicated significant enrichment of the ribosome and oxidative phosphorylation for the main effect of Bt at this time point, while the main effect of gregarines was once again enriched for ribosomes and coinfection for sphingolipid metabolism. GO terms most strongly associated with co-infection (cyan module) were primarily related to ribosomes, translation, and protein synthesis (**Table S15**). Other co-infection associated modules (darkgreen, green, lightcyan, and lightgreen) were enriched for chitin binding and various metabolic processes, although these did not survive FDR correction (**Table S15**). The eight-hour DE, KEGG, and co-expression results generally recapitulated the six-hour and gut results but had fewer modules with weaker associations, possibly due to the resolution of the acute immune response (**Tables S11, S13**).

### The net effect of gregarine infection is increased disease-induced mortality upon coinfection associated with a reduction in gut tolerance to damage

In size-matched larvae, gregarine infection significantly increased Bt-induced mortality relative to uninfected and well-fed larvae and exhibited no significant interaction effect with diet (Fig. 3A). To determine whether the extra mortality was due to differences in resistance among gregarine-infected and uninfected beetles, we measured Bt load via RT-qPCR across the 12 hour acute phase when most mortality is initiated (Fig. 3B). While bacterial load significantly increased and bifurcated over time as previously described in this and several other insect species (Duneau *et al*. 2017; Tate *et al*. 2017; Franz *et al*. 2023), gregarine samples were equally represented in high bacterial load trajectories (Bernoulli glm; z = 0.69, p = 0.49) relative to gregarine-uninfected samples, and neither gregarine exposure, confirmed gregarine infection, or the interaction of gregarines and time predicted bacterial load overall (Table 2, **Table S16**).

**Figure 3:**
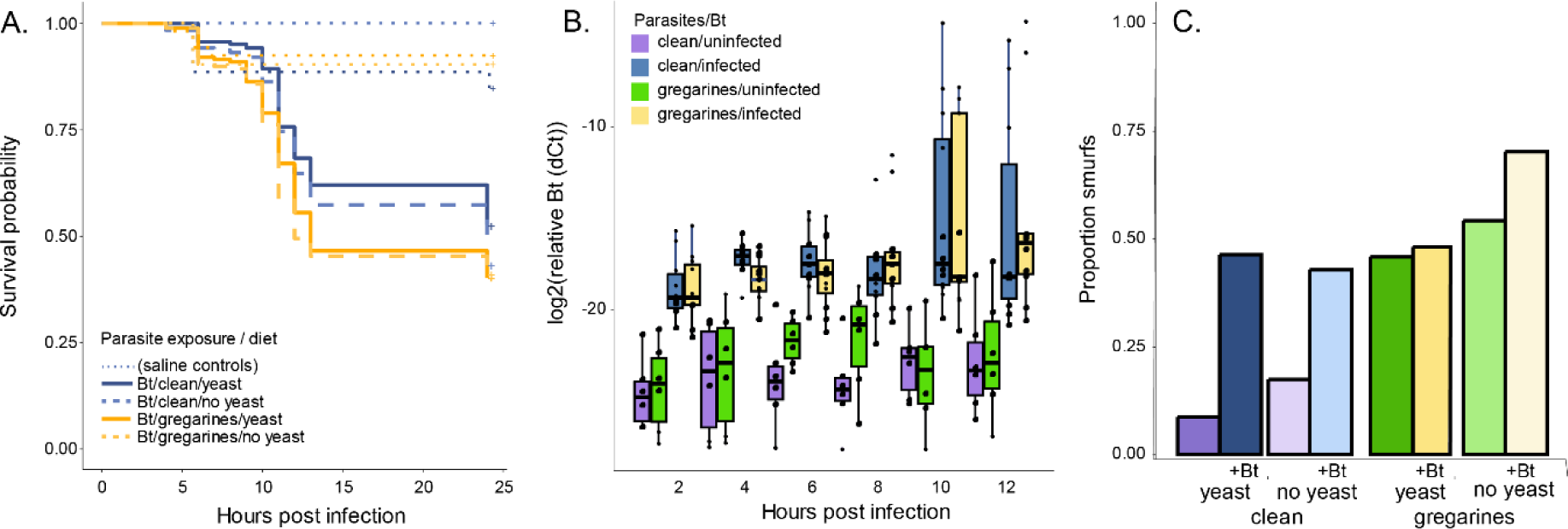
The impact of gregarines on host outcomes after Bt infection. A) Survival curves illustrate that saline-challenged larvae (dotted lines; same colors as Bt-infected legend) have high rates of survival regardless of treatment, but gregarine-infected larvae (yellow) are more likely to die than clean larvae (blue) after Bt infection regardless of diet (dashed: no yeast). B) Bacterial load relative to housekeeping gene (18s) was quantified via qPCR in Bt-infected (blue: no gregs, yellow: gregs) and saline control larvae (purple: no gregs, green: gregs), revealing no difference in Bt load based on gregarine status; all stats in Table SJJ. C) The proportion of larvae revealing smurf (leaky gut) phenotypes by gregarine, diet, and Bt infection treatment.

**Table 2.** The impact of coinfection on survival, Bt resistance, and gut integrity.

| *Survival post Bt or saline challenge*<sup>1</sup> |  |  |  |  |
| --- | --- | --- | --- | --- |
| Factor | exp(coef) | se(coef) | z | p |
| Bt challenge | 9.47 | 0.25 | 9.05 | **<0.0001** |
| Gregarine exposure | 1.39 | 0.14 | 2.39 | **0.017** |
| No-yeast diet | 1.15 | 0.14 | 0.98 | 0.32 |
| Gregarines:no-yeast | 0.93 | 0.19 | -0.39 | 0.7 |
| *Bacterial load over time by gregarine exposure status*<sup>2</sup> |  |  |  |  |
|  | estimate | se | t | p |
| Intercept | -19.30 | 1.04 | -18.57 | **< 0.001** |
| Time post Bt challenge | 0.33 | 0.13 | 2.56 | **0.012** |
| Gregarine exposure | -0.77 | 1.42 | -0.54 | 0.59 |
| Time:gregarines | 0.08 | 0.18 | 0.45 | 0.65 |
| *Bacterial load over time by confirmed greg infection status*<sup>2</sup> |  |  |  |  |
| Intercept | -19.43 | 0.83 | -23.32 | **< 0.001** |
| Time post Bt challenge | 0.34 | 0.11 | 3.12 | **0.0023** |
| Gregarine infection | -1.02 | 1.56 | -0.66 | 0.51 |
| Time:gregarines | 0.15 | 0.20 | 0.76 | 0.45 |
| *Proportion smurfs by gregarine exposure, Bt infection, and diet*<sup>3</sup> |  |  |  |  |
|  | estimate | se | z | p |
| Intercept | -2.18 | 0.75 | -2.90 | **0.0037** |
| Experimental block | -0.39 | 0.31 | -1.28 | 0.20 |
| Gregarine exposure | 2.21 | 0.85 | 2.60 | **0.0093** |
| Bt challenge | 2.23 | 0.83 | 2.68 | **0.0074** |
| No-yeast diet | 0.80 | 0.92 | 0.86 | 0.39 |
| Gregarines:Bt | -2.14 | 1.01 | -2.13 | **0.033** |
| Gregarines:diet | -0.46 | 1.09 | -0.42 | 0.67 |
| Bt:diet | -0.93 | 1.07 | -0.87 | 0.39 |
| Gregarines:Bt:Diet | 1.54 | 1.35 | 1.14 | 0.25 |

Notes: 1. Full Cox proportional hazards model is survival ~ Bt + gregarines\*diet + (1|block). Exp(coef) is the hazard ratio. Stratifying by bacterial vs saline challenge did not qualitatively change results. 2. Full linear model is log2(relative Bt load) ~ time\*gregarines. Includes only those larvae challenged with Bt; background amplification rates in controls were not significantly impacted by any factors (Table SJJ). Gregarine infection was confirmed by qPCR. 3. Full logistic model is smurf status ~ block +gregarines\*diet\*Bt. Reduced models did not yield a significantly better fit so the full factorial model is presented here.

Thus, gregarine-infected individuals are not less resistant to Bt. To understand whether tolerance mechanisms might instead account for the difference in mortality, we employed a smurf assay, which indicates gut leakiness through failure to maintain gut integrity or repair damaged structures (Fig. 3C). We found that both gregarine infection (Table 2) z = 2.6, p = 0.0093) and Bt infection (z = 2.7, p = 0.0074) individually predicted greater gut leakiness. Bt infection modestly increased smurf outcomes in coinfected beetles but not as drastically as in gregarine-free beetles (z = -2.1, p = 0.033). Neither diet alone nor its interaction terms contributed significantly to smurf status.

## Discussion

To what extent does a primary infection alter the metabolic and immunological landscape encountered by a second parasite species, and does resource allocation ultimately drive these differences? These questions are critical for building generalizable frameworks to predict the consequences of coinfection in natural populations and at different levels of biological organization. By manipulating resource availability and monitoring both metabolic and immunological facets of the host response to coinfection, we tested the resource sensitivity of key infection outcome parameters in a model system. Our results suggest that host development time, which contributes to age-structured infection susceptibility (Clay *et al*. 2023) and population intrinsic growth rate (Pearl *et al*. 1941; Park 1948), is exacerbated by the dual effects of gregarines and resource limitation. On the other hand, disease-induced mortality, which influences epidemiological dynamics and competitive outcomes in coinfected assemblages (Cortez & Duffy 2020; Rovenolt & Tate 2022), is not as sensitive to resources; differences are instead attributable to immune-related pathology during coinfection. Thus, mechanistic models that rely primarily on metabolic theory or energy budgets to predict coinfection dynamics are likely to underestimate the contribution of immunological shifts.

In our study, both resource quality and gregarine infection affected development time, presumably through slower storage of resources needed to grow. After accounting for development rate (*i.e.* with mass- rather than age-matched larvae), the metabolic state of diet-restricted and well-fed but gregarine-infected larvae largely catches up to their reference peers, leaving only the dual-stressed group with major metabolic consequences. This indicates that gregarines are capable of starving their hosts or forcing them to purge metabolites to avoid oxidative stress (Li *et al*. 2020), but the effects are dramatic only under resource-limited conditions; otherwise, the larvae appear to compensate by feeding more over a longer developmental window. This result aligns with our general understanding of gregarine infections as ubiquitous but relatively benign resource-exploiting parasites that inflict noticeable costs to their insect hosts only under multiple stressors or high parasite burdens (Randall *et al*. 2013; Wolz *et al*. 2022), and sets the stage for the phenotypes we observe upon Bt infection.

The mRNA-seq data suggest that gregarine infection alters the immune environment in the gut through the differential regulation of antimicrobial peptides and other effectors. This largely concurs with our previous study on gut gene expression after gregarine infection in the related flour beetle *T. castaneum*, although the latter exhibited much broader downregulation of antibacterial genes (Critchlow *et al*. 2019). This raises an interesting hypothesis that gregarines may differentially affect susceptibility to coinfection in these two co-occurring and competing host species (Park 1948; Rovenolt & Tate 2022). The transcriptional data also suggest that the gregarines affect the gut metabolic environment more generally, which may be important for nutrient processing and damage repair. The smurf assay indicates that diet does not significantly affect gut integrity, whereas gregarine-infected individuals have a greater baseline ‘leakiness’ regardless of diet or coinfection status. This is clearly not enough to kill them in isolation, since naïve and saline-stabbed larvae have low mortality rates regardless of their gregarine status (Fig. 3A). Bt infection initiates significant damage to the guts (also shown in (Critchlow *et al*. 2024)), but once everyone is infected Bt, the impact of gregarine status on additional gut leakiness is greatly diminished (Fig. 3C). Thus, there must be another contributor of pathology in gregarine- infected individuals to explain the difference in Bt-induced mortality.

Is it the altered immune environment? One set of co-expressed AMPs is specific to gregarine infection (as determined by gut expression) while another set is induced by Bt and not gregarines in the whole body. It is interesting that at the whole-body level, the first set is highly expressed specifically upon coinfection (rather than gregarines alone), suggesting that coinfection activates a separate immune program in the fat body and other tissues beyond the gut. Moving beyond specific AMPs, the WGCNA modules significantly associated with Bt infection feature a lot of the same players previously identified in RNA-seq studies of Bt in flour beetles (e.g. bacterial recognition, immune signaling and defense molecules, cytochrome P450s, serine proteases, glycolysis enzymes (Behrens *et al*. 2014; Tate & Graham 2017)). The modules most significantly associated with coinfection, however, are full of protein synthesis and metabolic genes, suggesting a struggle to effectively manage the physiological response.

Interestingly, there is not an observable difference in bacterial load between gregarine-infected and uninfected larvae, suggesting that neither mortality differences nor gene expression patterns are due to differences in resistance. Instead, these coinfection-specific modules point to differences in infection tolerance (Louie *et al*. 2016), possibly due to increased pathology of the alternate immune responses and/or their co-expressed genes or an increased struggle to maintain homeostatic metabolic and tissue repair programs. As we further improve the annotation of the *T. confusum* genome, we will be able to test these hypotheses with functional genomics approaches.

While we did not directly measure the production and spread of transmission stages, our results do hint at the consequences of coinfection for parasite fitness. Bt is an obligate killer, and relies on making spores in its dying or newly dead host to achieve transmission (Garbutt *et al*. 2011). Gregarines, on the other hand, mate in the living host gut to produce oocysts that are shed into the environment, and a dying host also spells a dead end for gregarines (Janovy *et al*. 2007). Thus, the exacerbated mortality in coinfected hosts undoubtedly hurts gregarine fitness, but it is not entirely clear that Bt benefits because Bt loads were not higher in coinfected individuals at the time of peak mortality. Future studies would benefit from new protocols for accurately quantifying gregarine transmission so that we can understand how this class of parasites, ubiquitous in the arthropod world (Rueckert *et al*. 2019), influence disease dynamics for biopesticides and vectored infections that preoccupy agricultural and biomedical efforts.

In conclusion, resources clearly matter – both the mRNA-seq and phenotype data suggest that the gregarines are indeed acting like parasites in depriving their hosts of resources and altering metabolic efficiency. Based on the metabolite data, the host can compensate for the parasitism when resources aren’t strictly limiting, but gregarine presence does change the immunological landscape in the face of secondary infection and may exert additional pathological effects. When it comes to infection mortality outcomes, the shifting immune landscape and physical damage inflicted by the gut parasite overshadow the importance of variance in resources. Thus, mechanistic models should allow for resource-independent contributions of immune responses when predicting or generalizing coinfection dynamics.

## Acknowledgements

We thank Justin Buchanan and James Deng for optimizing metabolic assay protocols for *Tribolium*, Jakob Heiser for assistance with the smurf assay, and Jacob Steenwyk for assistance with cd-hit. The experiments in this study were funded by NSF award 1753982 to A.T.T.; the *T. confusum* transcriptome assembly was funded by NIH award R35GM138007 to A.T.T.

## Data Availability

RNA-seq data is publicly available in the NCBI Sequence Read Archive (project accession PRJNA771764). Experimental data will be available in Data Dryad (DOI in progress).

## Supplementary Tables

Table S1 Sample identity and replication for mRNA-seq

Table S2 Comprehensive RNA sample information (replicate number, sample description, file size, bowtie statistics)

Table S3 Contig statistics

Table S4 Busco results

Table S5 Training sets used to identify AMPs via HMM

Table S6 Differential expression analysis for the gut upon infection with gregarine parasites

Table S7 AMP names and expression patterns

Table S8 WGCNA analysis results for the gut

Table S9 KEGG analysis on differentially expressed genes

Table S10 Differential expression analysis for the whole body 6 hours post-infection

Table S11 Differential expression analysis for the whole body 8 hours post-infection

Table S12 WGCNA analysis for the whole body 6 hours post-infection

Table S13 WGCNA analysis for the whole body 8 hours post-infection

Table S14 Modules of co-expressed AMPs using WGCNA on whole body samples

Table S15 GO analyses for modules significantly associated with Bt or co-infection

Table S16 Extended statistics for survival and bacterial load assays

## Supplementary Methods and Figures

Figure S1. Bit score values for BlastP and BlastX results.

Figure S2. Expression of effector genes including antimicrobial peptides (AMPs) and pathogenesis-related (PR) proteins.

Figure S3. Relationship between individual larval mass and mass-normalized metabolite levels

Figure S4. Principal component analyses (PCAs) for three differential expression analyses.

Figure S5. The posterior probabilities of differential expression using Bayseq.

Figure S6. WGCNA analyses for the whole body 6 and 8 hours post-infection.

## References

1. Adamo, S.A., Davies, G., Easy, R., Kovalko, I. & Turnbull, K.F. (2016). Reconfiguration of the immune system network during food limitation in the caterpillar Manduca sexta. The Journal of Experimental Biology.

2. Andrews, S. (2010). FastQC: A quality control tool for high throughput sequence data. Reference Source.

3. Barr, J.S., Estevez-Lao, T.Y., Khalif, M., Saksena, S., Yarlagadda, S., Farah, O. et al. (2023). Temperature and age, individually and interactively, shape the size, weight, and body composition of adult female mosquitoes. Journal of Insect Physiology, 148, 104525.

4. Bates D., M.M. (2010). lme4: linear mixed-effects models using S4 classes.

5. Behrens, S., Peuss, R., Milutinovi, B., Eggert, H., Esser, D., Rosenstiel, P. et al. (2014). Infection routes matter in population-specific responses of the red flour beetle to the entomopathogen Bacillus thuringiensis. BMC Genomics, 15, 445.

6. Benjamini, Y. & Hochberg, Y. (1995). Controlling the false discovery rate: a practical and powerful approach to multiple testing. Journal of the Royal statistical society: series B (Methodological), 57, 289–300.

7. Bernot, R.J. & Poulin, R. (2018). Ecological Stoichiometry for Parasitologists. Trends in Parasitology, 34, 928–933.

8. Bray, N.L., Pimentel, H., Melsted, P. & Pachter, L. (2016). Near-optimal probabilistic RNA-seq quantification. Nature biotechnology, 34, 525–527.

9. Clay, P.A., Gattis, S., Garcia, J., Hernandez, V., Ben-Ami, F. & Duffy, M.A. (2023). Age Structure Eliminates the Impact of Coinfection on Epidemic Dynamics in a Freshwater Zooplankton System. The American Naturalist, 202, 785–799.

10. Cortez, M.H. & Duffy, M.A. (2020). Comparing the Indirect Effects between Exploiters in Predator-Prey and Host-Pathogen Systems. The American Naturalist, 196, E144–E159.

11. Cressler, C.E., Nelson, W.A., Day, T. & McCauley, E. (2014). Disentangling the interaction among host resources, the immune system and pathogens. Ecology Letters, 17, 284–293.

12. Critchlow, J.T., Norris, A. & Tate, A.T. (2019). The legacy of larval infection on immunological dynamics over metamorphosis. Philosophical Transactions of the Royal Society B: Biological Sciences, 374, 20190066.

13. Critchlow, J.T., Prakash, A., Zhong, K.Y. & Tate, A.T. (2024). Mapping the functional form of the trade-off between infection resistance and reproductive fitness under dysregulated immune signaling. PLOS Pathogens, 20, e1012049.

14. Deschodt, P.S. & Cory, J.S. (2022). Resource limitation has a limited impact on the outcome of virus–fungus co-infection in an insect host. Ecology and Evolution, 12, e8707.

15. Detwiler, J. & Janovy, J., Jr. (2008). The role of phylogeny and ecology in experimental host specificity: Insights from a eugregarine-host system. Journal of Parasitology, 94, 7–12.

16. Duneau, D., Ferdy, J.-B., Revah, J., Kondolf, H., Ortiz, G.A., Lazzaro, B.P. et al. (2017). Stochastic variation in the initial phase of bacterial infection predicts the probability of survival in D. melanogaster. eLife, 6, e28298.

17. Finn, R.D., Clements, J. & Eddy, S.R. (2011). HMMER web server: interactive sequence similarity searching. Nucleic acids research, 39, W29–W37.

18. Franz, M., Armitage, S.A., Rolff, J. & Regoes, R.R. (2023). Virulence decomposition for bifurcating infections. Proceedings of the Royal Society B, 290, 20230396.

19. Fu, L., Niu, B., Zhu, Z., Wu, S. & Li, W. (2012). CD-HIT: accelerated for clustering the next- generation sequencing data. Bioinformatics, 28, 3150–3152.

20. Garbutt, J., Bonsall, M.B., Wright, D.J. & Raymond, B. (2011). Antagonistic competition moderates virulence in Bacillus thuringiensis. Ecology Letters, 14, 765–772.

21. Grabherr, M.G., Haas, B.J., Yassour, M., Levin, J.Z., Thompson, D.A., Amit, I. et al. (2011). Full-length transcriptome assembly from RNA-Seq data without a reference genome. Nature biotechnology, 29, 644–652.

22. Graham, A.L. (2008). Ecological rules governing helminth–microparasite coinfection. Proceedings of the National Academy of Sciences, 105, 566–570.

23. Griffiths, E.C., Fairlie-Clarke, K., Allen, J.E., Metcalf, C.J.E. & Graham, A.L. (2015). Bottom- up regulation of malaria population dynamics in mice co-infected with lung-migratory nematodes. Ecology Letters, 18, 1387–1396.

24. Hardcastle, T.J. & Kelly, K.A. (2010). baySeq: empirical Bayesian methods for identifying differential expression in sequence count data. BMC bioinformatics, 11, 1–14.

25. Herndon, N., Shelton, J., Gerischer, L., Ioannidis, P., Ninova, M., Dönitz, J. et al. (2020). Enhanced genome assembly and a new official gene set for Tribolium castaneum. BMC Genomics, 21, 47.

26. Huang, D.W., Sherman, B.T., Tan, Q., Collins, J.R., Alvord, W.G., Roayaei, J. et al. (2007). The DAVID Gene Functional Classification Tool: a novel biological module-centric algorithm to functionally analyze large gene lists. Genome biology, 8, 1–16.

27. Janovy, J., Jr., Detwiler, J., Schwank, S., Bolek, M.G., Knipes, A.K. & Langford, G.J. (2007). New and emended descriptions of gregarines from flour beetles (Tribolium spp. and Palorus subdepressus: Coleoptera, Tenebrionidae). Journal of Parasitology, 93, 1155- 1170.

28. Jent, D., Perry, A., Critchlow, J. & Tate, A.T. (2019). Natural variation in the contribution of microbial density to inducible immune dynamics. Molecular Ecology, 28, 5360–5372.

29. Johnson, P.T.J., de Roode, J.C. & Fenton, A. (2015). Why infectious disease research needs community ecology. Science, 349, 1259504.

30. Langfelder, P. & Horvath, S. (2008). WGCNA: an R package for weighted correlation network analysis. BMC Bioinformatics, 9, 559.

31. Li, X., Rommelaere, S., Kondo, S. & Lemaitre, B. (2020). Renal Purge of Hemolymphatic Lipids Prevents the Accumulation of ROS-Induced Inflammatory Oxidized Lipids and Protects Drosophila from Tissue Damage. Immunity, 52, 374–387.e376.

32. Louie, A., Song, K.H., Hotson, A., Thomas Tate, A. & Schneider, D.S. (2016). How Many Parameters Does It Take to Describe Disease Tolerance? PLoS Biol, 14, e1002435.

33. Love, M.I., Huber, W. & Anders, S. (2014). Moderated estimation of fold change and dispersion for RNA-seq data with DESeq2. Genome Biology, 15, 550.

34. Manni, M., Berkeley, M.R., Seppey, M., Simão, F.A. & Zdobnov, E.M. (2021). BUSCO update: novel and streamlined workflows along with broader and deeper phylogenetic coverage for scoring of eukaryotic, prokaryotic, and viral genomes. Molecular biology and evolution, 38, 4647–4654.

35. Ott, D., Digel, C., Rall, B.C., Maraun, M., Scheu, S. & Brose, U. (2014). Unifying elemental stoichiometry and metabolic theory in predicting species abundances. Ecology Letters, 17, 1247–1256.

36. Park, T. (1948). Interspecies Competition in Populations of Trilobium confusum Duval and Trilobium castaneum Herbst. Ecological Monographs, 18, 265–307.

37. Park, T., Nathanson, M., Ziegler, J.R. & Mertz, D.B. (1970). Cannibalism of Pupae by Mixed- Species Populations of Adult Tribolium. Physiological Zoology, 43, 166–184.

38. Pearl, R., Park, T. & Miner, J.R. (1941). Experimental Studies on the Duration of Life. XVI. Life Tables for the Flour Beetle Tribolium confusum Duval. The American Naturalist, 75, 5–19.

39. R_Core_Team (2012b). R: a language and environment for statistical computing. In: Vienna, *Austria*: *R foundation for statistical computing*.

40. Ramesh, A. & Hall, S.R. (2023). Niche theory for within-host parasite dynamics: Analogies to food web modules via feedback loops. Ecology Letters, 26, 351–368.

41. Randall, J., Cable, J., Guschina, I.A., Harwood, J.L. & Lello, J. (2013). Endemic infection reduces transmission potential of an epidemic parasite during co-infection. Proceedings of the Royal Society of London B: Biological Sciences, 280, 20131500.

42. Rovenolt, F. & Tate, A.T. (2022). The impact of coinfection dynamics on host competition and coexistence. The American Naturalist, 199, 91–107.

43. Rueckert, S., Betts, E.L. & Tsaousis, A.D. (2019). The Symbiotic Spectrum: Where Do the Gregarines Fit? Trends in Parasitology, 35, 687–694.

44. Rynkiewicz, E.C., Pedersen, A.B. & Fenton, A. (2015). An ecosystem approach to understanding and managing within-host parasite community dynamics. Trends in Parasitology, 31, 212–221.

45. Schulz, N.K.E., Stewart, C.M. & Tate, A.T. (2023). Female investment in terminal reproduction or somatic maintenance depends on infection dose. Ecological Entomology, 48, 714–724.

46. Seguel, M., Budischak, S.A., Jolles, A.E. & Ezenwa, V.O. (2023). Helminth-associated changes in host immune phenotype connect top-down and bottom-up interactions during co- infection. Functional Ecology, 37, 860–872.

47. Tate, A.T. (2019). Role of Multiple Infections on Immunological Variation in Wild Populations. mSystems, 4, e00099–00019.

48. Tate, A.T., Andolfatto, P., Demuth, J.P. & Graham, A.L. (2017). The within-host dynamics of infection in trans-generationally primed flour beetles. Molecular Ecology, 26, 3794– 3807.

49. Tate, A.T. & Graham, A.L. (2015). Trans-generational priming of resistance in wild flour beetles reflects the primed phenotypes of laboratory populations and is inhibited by co-infection with a common parasite. Functional Ecology, 29, 1059–1069.

50. Tate, A.T. & Graham, A.L. (2017). Dissecting the contributions of time and microbe density to variation in immune gene expression. Proceedings of the Royal Society B: Biological Sciences, 284.

51. Therneau, T. (2014). A Package for Survival Analysis in S. R package version 2.37–7. 2014.

52. Therneau, T.M. & Therneau, M.T.M. (2015). Package ‘coxme’. *R package version*, 2.

53. Thomas, A.M. & Rudolf, V.H. (2010). Challenges of metamorphosis in invertebrate hosts: Maintaining parasite resistance across life-history stages. Ecological Entomology, 35, 200–205.

54. Vale, P., F., Choisy, M. & Little Tom, J. (2013). Host nutrition alters the variance in parasite transmission potential. Biology Letters, 9, 20121145.

55. Wolz, M., Rueckert, S. & Müller, C. (2022). Fluctuating Starvation Conditions Modify Host-Symbiont Relationship Between a Leaf Beetle and Its Newly Identified Gregarine Species. Frontiers in Ecology and Evolution, 10.

56. Yu, G., Wang, L.-G., Han, Y. & He, Q.-Y. (2012). clusterProfiler: an R package for comparing biological themes among gene clusters. Omics: a journal of integrative biology, 16, 284–287.

57. Zanchi, C., Lindeza, A.S. & Kurtz, J. (2020). Comparative mortality and adaptation of a smurf assay in two species of tenebrionid beetles exposed to Bacillus thuringiensis. Insects, 11, 261.

58. Zilio, G. & Koella, J.C. (2020). Sequential co-infections drive parasite competition and the outcome of infection. Journal of Animal Ecology, 89, 2367–2377.

