## Supplemental Methods and Figures for "Resources modulate developmental shifts but not infection tolerance upon coinfection in an insect system"

#### RNA samples

We used the RNA samples listed in Table S2 for RNA sequencing.

#### *T. confusum* transcriptome assembly

We first assessed RNAseq read quality using fastqc (Andrews 2010). Using only samples not infected with gregarines, we assembled a *de novo* transcriptome using Trinity with default settings; quality filtering was performed within Trinity and reads were assembled in paired-end mode (Grabherr et al. 2011). Highly similar transcripts were clustered using cd-hit (Fu et al. 2012). To assess the quality of the assembly, the reads were realigned to the assembled transcriptome using bowtie2 and the ExN50 statistic was calculated within Trinity (Grabherr et al. 2011) (Table S3). To assess the completeness of the assembly, transcripts were analyzed using BUSCO (v.5.2.2) (Benchmarking Universal Single-Copy Orthologs) against an insect gene set (Manni et al. 2021) (Table S4).

#### Annotating the *T. confusum* transcriptome

We annotated the *T. confusum* transcriptome by using tblastx with parameters *-evalue 0.01*, *-max_target_seqs 1*, and *-max_hsps 1* to blast the transcriptome to the cDNA reference of *T. castaneum* (GCA_000002335.3). Next, the isoform numbers were removed from the ends of both the query sequences and the database sequences, leaving just the gene IDs. We then removed duplicate hits from the blast output file. After this, there were 77,812 total entries in the file, representing 72,241 unique *T. confusum* sequences (defined as “genes”) and 13,892 *T. castaneum* genes (from the reference transcriptome). From here, we did a final collapsing step; we chose the best blast hit for each *T. confusum* sequence (based on lowest e-value) to collapse to 72,241 entries, where each each *T. confusum* sequence was represented once.

#### Identifying gregarine contaminants

To identify contaminating reads, we used blastn with parameters *-evalue 0.01*, *-max_target_seqs 1*, and *-max_hsps 1* to blast the transcriptome against the *Gregarina niphandrodes* cDNA reference (GCA_000223845.4). This search produced hits for 79 Trinity isoforms, representing 34 unique “gene” names. E-values ranged from 0 to 4.5e-06.

#### AMP annotation

We used HMMER (Finn, Clements, and Eddy 2011) to annotate AMPs in the *T. confusum* proteome using Hidden Markov Models (HMMs). To this end, we used Coleoptera AMPs as training sets and constructed HMM profiles. We made three distinct training sets for glycine-rich peptides (Attacin/Diptericin), Cecropins, and Defensins. We downloaded training sets from the NCBI protein database using three keywords: “Coleoptera Attacin”, “Coleoptera Cecropin”, and “Coleoptera Defensin”. From the Attacin dataset, we removed a sequence with the ID: KAF5271830.1, because it poorly aligned with sequences in the Attacin set. In the Defensin dataset, five sequences were longer than 150 amino acids and thus we removed them to improve alignment. This left us with 105 Attacins, 83 Defensins, and 17 Cecropins (Supplemental table SR). AMP sequences in the three training datasets are deposited here: https://github.com/danialasg74/coleoptera-AMPs-training-sets-. We constructed HMM profiles using each training set. Next, we used HMM profiles to identify Antimicrobial Peptides (AMPs) within the proteome of *T. confusum.* We generated the proteome by identifying and translating the longest open reading frame for each transcript within the transcriptome.

#### Finding orthologs in *T. castaneum*

We identified orthologous *T. confusum* genes in *T. castaneum.* To this end, we blasted protein and transcript sequences of *T. confusum* against the protein database of *T. castaneum* using BlastP and Blatx respectively. Next, to examine the quality of matches, we plotted the bit scores for both blasts (Figure S1). Based on the inspection of plots, we chose a minimum bit score cutoff of 50 for BlastP and 100 for Blastx. The cutoff values were chosen to exclude the high-density region of the two graphs, which is caused by a bulk of spurious matches with low bit scores. With these cutoff values, we found 30,543 matches using BlastP and 29,986 matches with Blastx. Out of the matches identified by Blastx, 91% were identical to those found by

BlastP. However, 4,334 matches found by Blastx could not be identified with BlastP; thus, they were added to the results obtained from BlastP. This makes a total of 34,877 matches between *T. confusum* and *T. castaneum* genes. Several genes in *T. confusum* matched with identical genes in *T. castaneum*. After removing redundancy, we ended up with a total of 13,865 orthologs between *T. confusum* and *T. castaneum* genes.

#### Differential expression analysis

We used kallisto v 0.48.0 to quantify gene expression (Bray et al. 2016). To this end, we performed pseudo alignment of RNA-seq reads to the assembled transcriptome of *T. confusum*. We measured gene expression by summing count or transcript per million (TPM) values across isoforms. We blasted AMP transcripts against the *T. castaneum* transcriptome (Blastn) and removed similar sequences to AMPs. This improved the quantification of AMP gene expression.

We used the DEseq2 (v 1.36) package in R (Love, Huber, and Anders 2014) to run three differential expression (DE) analyses (Supplemental table SD), excluding very low-expressed genes (counts < 10). In the first analysis, we identified DE genes in the gut upon Greg infection. We used sample 1 as the control and sample 6 as the treatment group (Table S1). For this analysis, we modeled gene expression as $E \sim G$ in DEseq2, where $G$ represents Greg infection and $E$ is the expression level of each gene. In the second and third analyses, we identified DE genes upon infection with Bt, Greg, and co-infection by both Bt and Greg in the whole body samples. In both analyses, we used sample 2 as the control and sample 7 to evaluate the effect of Greg infection on gene expression. We chose these samples because they include a large number of replicates (eight replicates), which increased statistical power for the detection of DE genes. For the second analysis, we measured the effect of Bt and co-infection on gene expression six hours post-infection (groups 4 and 9), while in the third analysis, we measured the effect of Bt and co-infection eight hours post-infection (groups 5 and 10). For both analyses, we measured the effect of Greg at $t = 0$. We modeled gene expression in DEseq2 as $E \sim B + G + B \times G$. Here $B$ is the effect of Bt, $G$ is the effect of Greg, and $B \times G$ is the effect of co-infection. For the second and third analyses, we also used the Bayseq (2.30) package in R (Hardcastle and Kelly 2010). To this end, we divided samples into four groups: genes that are not DE across samples, genes that are DE in samples infected by Bt relative to other samples, genes that are DE in the Greg infected samples relative to other samples, and genes that are DE in co-infected samples relative to other samples.

For DEseq analyses, we considered a gene to be differentially expressed if the false-discovery rate (FDR) corrected *P*-value was below 0.05 (Benjamini and Hochberg 1995). For analyses performed via Bayseq, we reported the posterior probability of differential expression.

#### Co-expression network analysis

We performed weighted gene co-expression network analysis (WGCNA) to identify modules of co-expressed genes. We conducted three WGCNA analyses on genes that have orthologs in *T. castaneum,* similar to the three DE analyses reported in Supplemental table SD. Genes with zero count values across all replicates were removed before analyses. Next, We constructed a signed correlation matrix for each analysis using the count data. We used signed correlation matrices to identify positive or negative correlations between the expression of genes in the network. To produce scale-free topology networks, we raised correlation matrices to the power of 20. Next, to identify clusters of genes with similar patterns of expression, we used a merging threshold of 0.25 and a minimum module size of 30. We calculated the Pearson correlation of the module eigengenes across samples to identify modules of co-expressed genes.

#### GO and KEGG analyses

We performed gene ontology (GO) analyses to find functional categories for co-expressed modules identified via WGCNA. Because GO terms were not available for *T. confusum,* we performed GO analyses on orthologous sequences in *T. castaneum.* We used DAVID Gene Functional Classification Tool for GO analysis (Huang et al. 2007).

We also performed Kyoto Encyclopedia of Genes and Genomes (KEGG) analyses on differentially expressed genes that were identified using DEseq2. To this end, we used the clusterProfiler package in R (Yu et al. 2012). We first made a list of genes with padj value less than 0.05 and log fold change greater than 1. Next, for these genes, we identified orthologous genes of *T. castaneum* on which we performed KEGG analyses.

### References:


1. Andrews S. (2010). FastQC: A Quality Control Tool for High Throughput Sequence Data [Online]. Available online at: http://www.bioinformatics.babraham.ac.uk/projects/fastqc/
2. Benjamini, Yoav, and Yosef Hochberg. 1995. “Controlling the False Discovery Rate: A Practical and Powerful Approach to Multiple Testing.” Journal of the Royal Statistical Society. Series B, Statistical Methodology 57 (1): 289–300.
3. Bray, Nicolas L., Harold Pimentel, Páll Melsted, and Lior Pachter. 2016. “Near-Optimal Probabilistic RNA-Seq Quantification.” Nat. Biotechnol. 34 (5): 525–27.
4. Finn, Robert D., Jody Clements, and Sean R. Eddy. 2011. “HMMER Web Server: Interactive Sequence Similarity Searching.” Nucleic Acids Research 39 (Web Server issue): W29–37.
5. Fu, Limin, Beifang Niu, Zhengwei Zhu, Sitao Wu, and Weizhong Li. 2012. “CD-HIT: Accelerated for Clustering the next-Generation Sequencing Data.” Bioinformatics 28 (23): 3150–52.
6. Grabherr, Manfred G., Brian J. Haas, Moran Yassour, Joshua Z. Levin, Dawn A. Thompson, Ido Amit, Xian Adiconis, et al. 2011. “Full-Length Transcriptome Assembly from RNA-Seq Data without a Reference Genome.” Nature Biotechnology 29 (7): 644–52.
7. Hardcastle, Thomas J., and Krystyna A. Kelly. 2010. “baySeq: Empirical Bayesian Methods for Identifying Differential Expression in Sequence Count Data.” BMC Bioinformatics 11 (August): 422.
8. Huang, Da Wei, Brad T. Sherman, Qina Tan, Jack R. Collins, W. Gregory Alvord, Jean Roayaei, Robert Stephens, Michael W. Baseler, H. Clifford Lane, and Richard A. Lempicki. 2007. “The DAVID Gene Functional Classification Tool: A Novel Biological Module-Centric Algorithm to Functionally Analyze Large Gene Lists.” Genome Biology 8 (9): R183.
9. Love, Michael I., Wolfgang Huber, and Simon Anders. 2014. “Moderated Estimation of Fold Change and Dispersion for RNA-Seq Data with DESeq2.” Genome Biol. 15 (12): 550.
10. Manni, Mosè, Matthew R. Berkeley, Mathieu Seppey, Felipe A. Simão, and Evgeny M. Zdobnov. 2021. “BUSCO Update: Novel and Streamlined Workflows along with Broader and Deeper Phylogenetic Coverage for Scoring of Eukaryotic, Prokaryotic, and Viral Genomes.” Molecular Biology and Evolution 38 (10): 4647–54.
11. Yu, Guangchuang, Li-Gen Wang, Yanyan Han, and Qing-Yu He. 2012. “clusterProfiler: An R Package for Comparing Biological Themes among Gene Clusters.” Omics: A Journal of Integrative Biology 16 (5): 284–87.

**Figure S1.** Bit score values plotted for BlastP and BlastX results. The x-axis is the bit scores and the y-axis is the frequency of bit scores. The red dashed line delineates the cutoff chosen to exclude poor matches with low bit scores.

**Figure S2.** Expression of effector genes including annotated antimicrobial peptides (AMPs) and pathogenesis-related (PR) proteins. The x-axis shows the samples and the y-axis shows the genes. The color bar shows the level of expression which is the log-transformed of normalized count data (TPM). Darker colors show lower levels of expression and brighter colors show higher expression.

**Figure S3**. Relationship between individual larval mass (in mg) and mass-normalized metabolite levels for metabolite assays in gregarine-unexposed (control) or exposed individuals in standard or no-yeast (low-protein) diets. Asterisks indicate a significant correlation.

**Figure S4.** Principal Component Analysis (PCA) plots show clustering of replicates (circles) for gregarine-infected (green), Bt-infected (blue) or corresponding uninfected (with the same parasite; purple) samples across three DE analyses. The three DE analyses are: infection of the gut with gregarines only or six and eight hours post infection with Bt in the whole body. The sample labels (AT numbers) are shown for each sample.

**Figure S5.** The posterior probabilities of differential expression using Bayseq relative to Deseq2 results. The y-axis shows the posterior probability for the effectors on the x-axis. For 6 hour whole body (first row) and 8 hours whole body, posterior probabilities are shown for three groups: co-infection, Bt infected, and greg infected. The blue circles are genes that are found to be significantly differentially expressed by Bt using DEseq2, while the green circles are genes that are found to be significantly differentially expressed by gregarines.

**Figure S6.** WGCNA analysis shows modules of co-expressed genes for whole body samples at six (top left) and eight hours (bottom left) post-infection. The red cells show a positive correlation, while blue cells show a negative correlation. The *P* values are reported beneath Pearson correlation magnitudes for different modules (color bar on the left) across samples (X-axis). The color bar on the right shows the magnitude of correlations. The two panels on the right show the expression of genes within each module as box plots across infection treatments at six or eight hours post Bt infection.

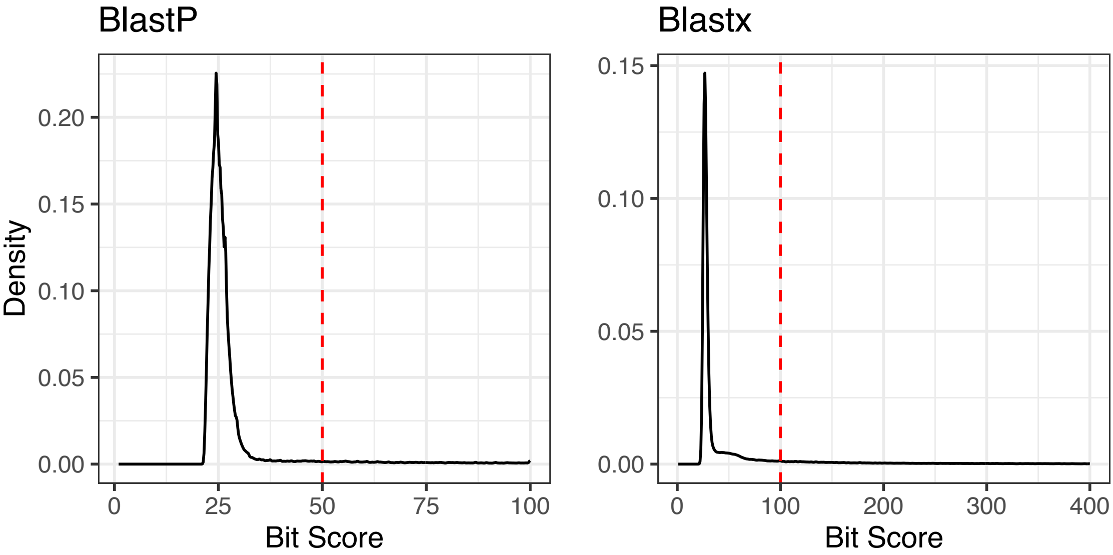
BlastP
Blastx
Density
0.20
0.15
0.10
0.05
0.00
0
25
50
75
100
Bit Score
0.15
0.10
0.05
0.00
0
100
200
300
400
Bit Score

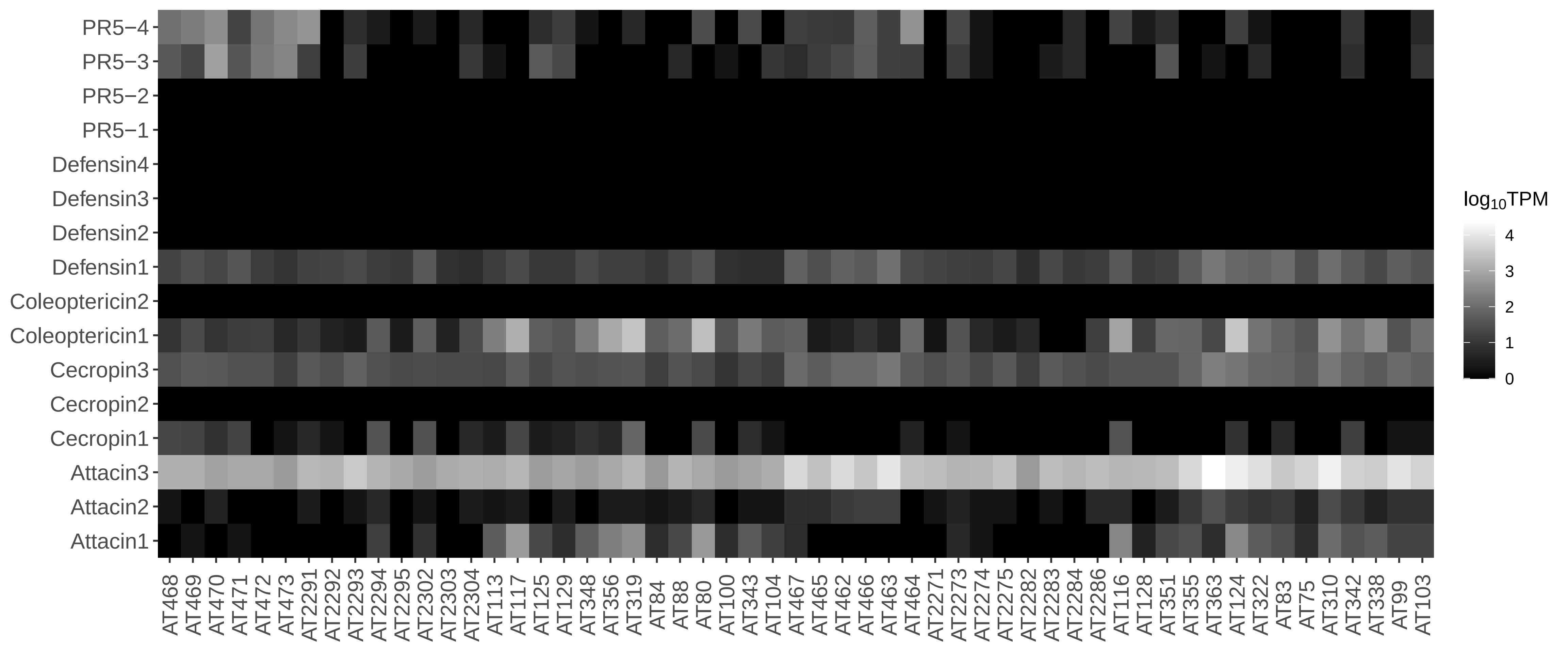

PR5−4
PR5−3
PR5−2
PR5−1
Defensin4
Defensin3
Defensin2
Defensin1
Coleoptericin2
Coleoptericin1
Cecropin3
Cecropin2
Cecropin1
Attacin3
Attacin2
Attacin1
AT468 AT469 AT470 AT471 AT472 AT473 AT2291 AT2292 AT2293 AT2294 AT2295 AT2302 AT2303 AT2304 AT113 AT117 AT125 AT129 AT348 AT356 AT319 AT84 AT88 AT80 AT100 AT343 AT104 AT467 AT465 AT462 AT466 AT463 AT464 AT2271 AT2273 AT2274 AT2275 AT2282 AT2283 AT2284 AT2286 AT116 AT128 AT351 AT355 AT363 AT124 AT322 AT83 AT75 AT310 AT342 AT338 AT99 AT103
$log_{10}$TPM
4
3
2
1
0

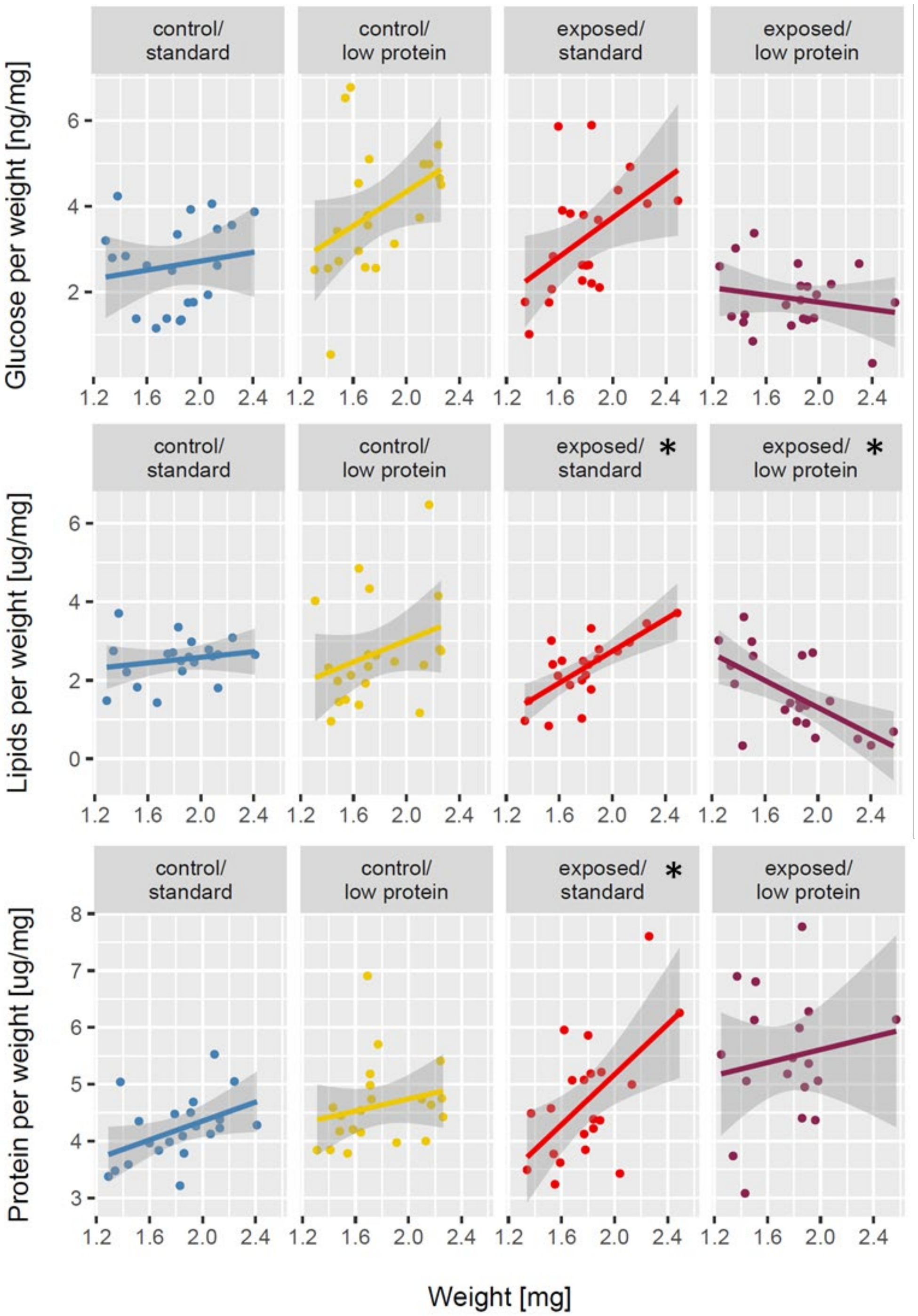


**Figure SFA**. Relationship between individual larval mass (in mg) and mass-normalized metabolite levels for metabolite assays in gregarine-unexposed (control) or exposed individuals in standard or no-yeast (low-protein) diets. Asterisks indicate a significant correlation.

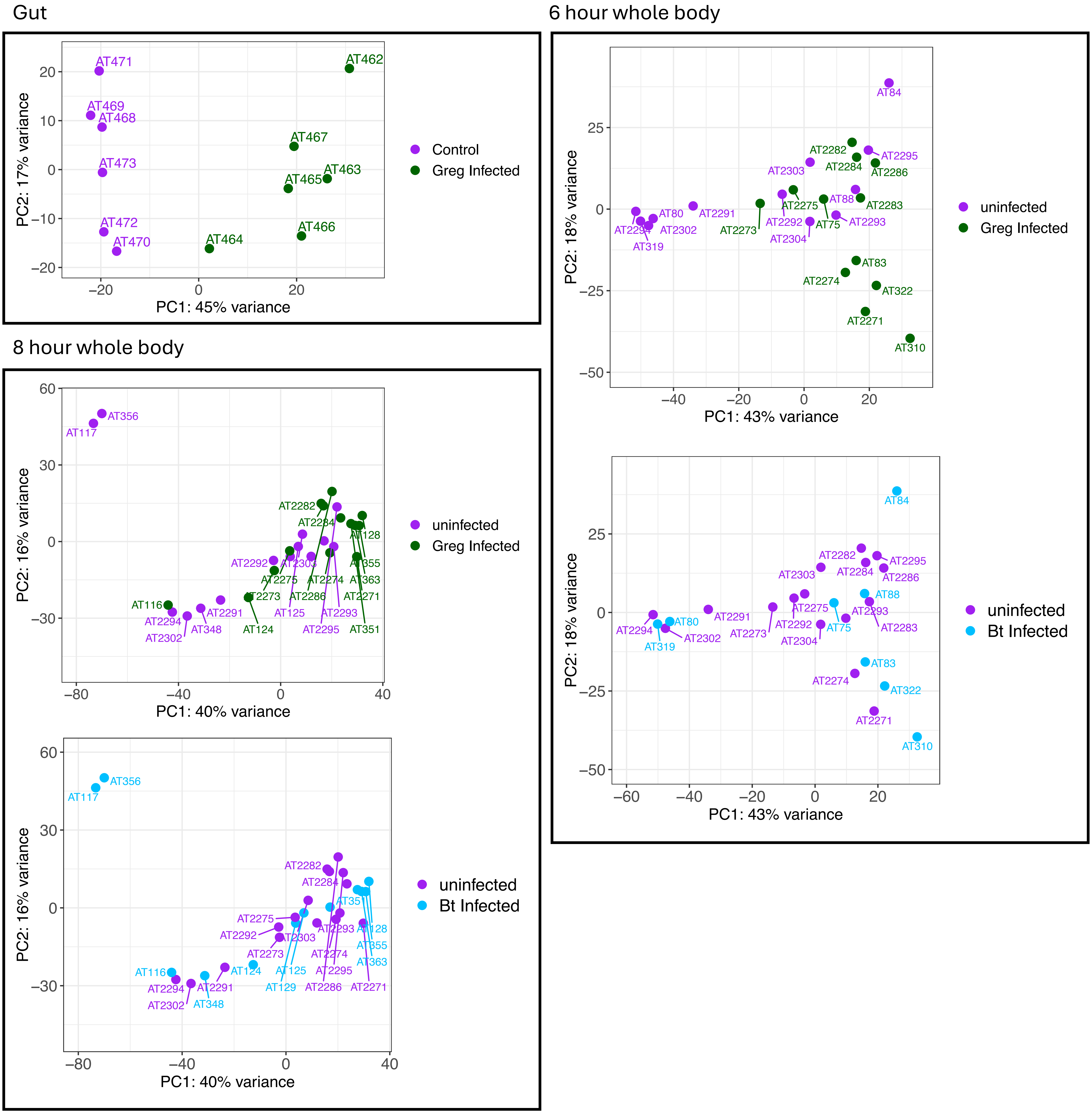
Gut
PC1: 45% variance
PC2: 17% variance
AT471
AT469
AT468
AT473
AT472
AT470
AT462
AT467
AT463
AT465
AT466
AT464
Control
Greg Infected
6 hour whole body
PC1: 43% variance
PC2: 18% variance
uninfected
Greg Infected
uninfected
Bt Infected
8 hour whole body
PC1: 40% variance
PC2: 16% variance
uninfected
Greg Infected
uninfected
Bt Infected

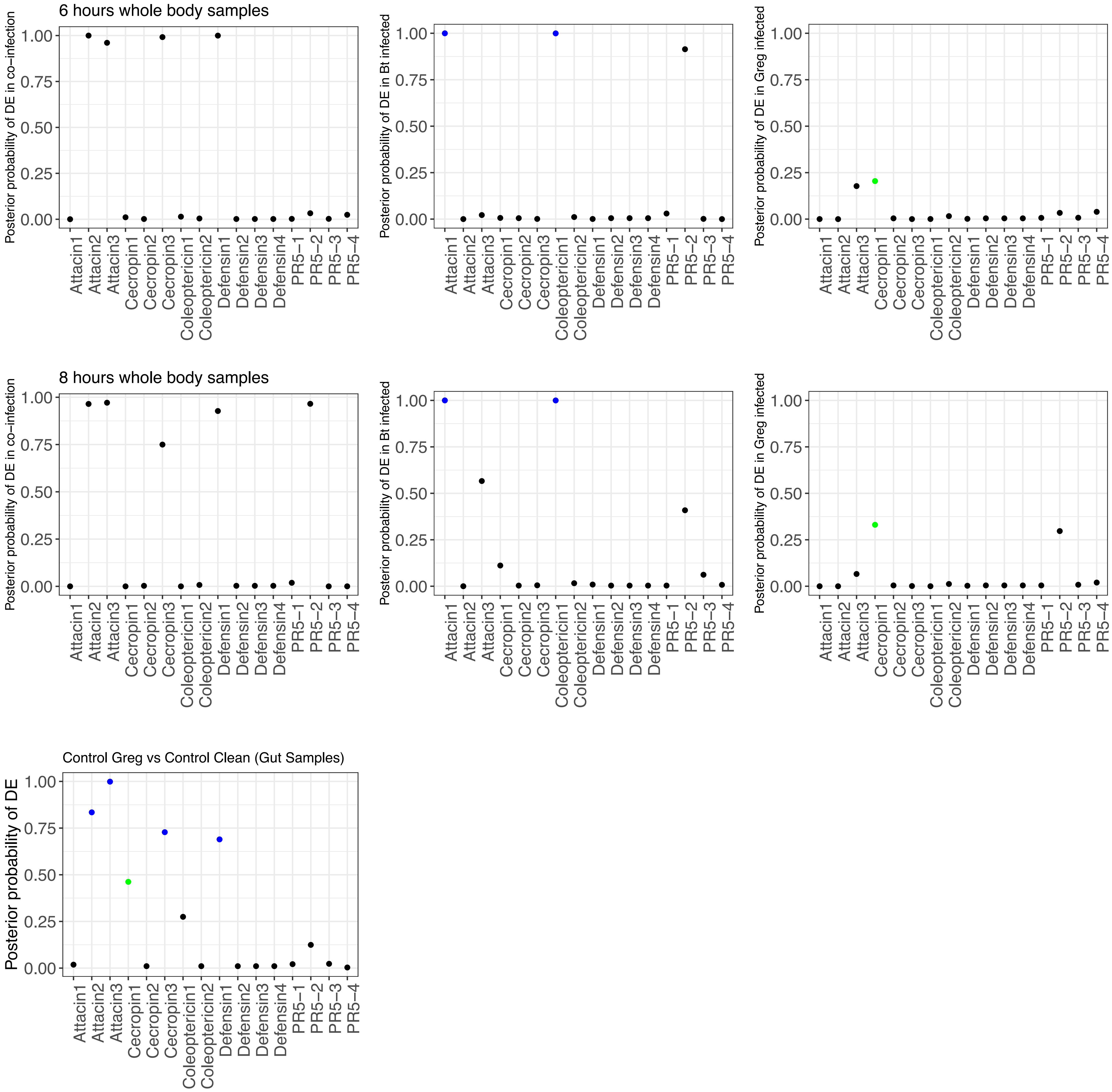
6 hours whole body samples
8 hours whole body samples
Control Greg vs Control Clean (Gut Samples)
Posterior probability of DE in co−infection
Posterior probability of DE in Bt infected
Posterior probability of DE in Greg infected
Posterior probability of DE
0.00
0.25
0.50
0.75
1.00
Attacin1
Attacin2
Attacin3
Cecropin1
Cecropin2
Cecropin3
Coleoptericin1
Coleoptericin2
Defensin1
Defensin2
Defensin3
Defensin4
PR5−1
PR5−2
PR5−3
PR5−4

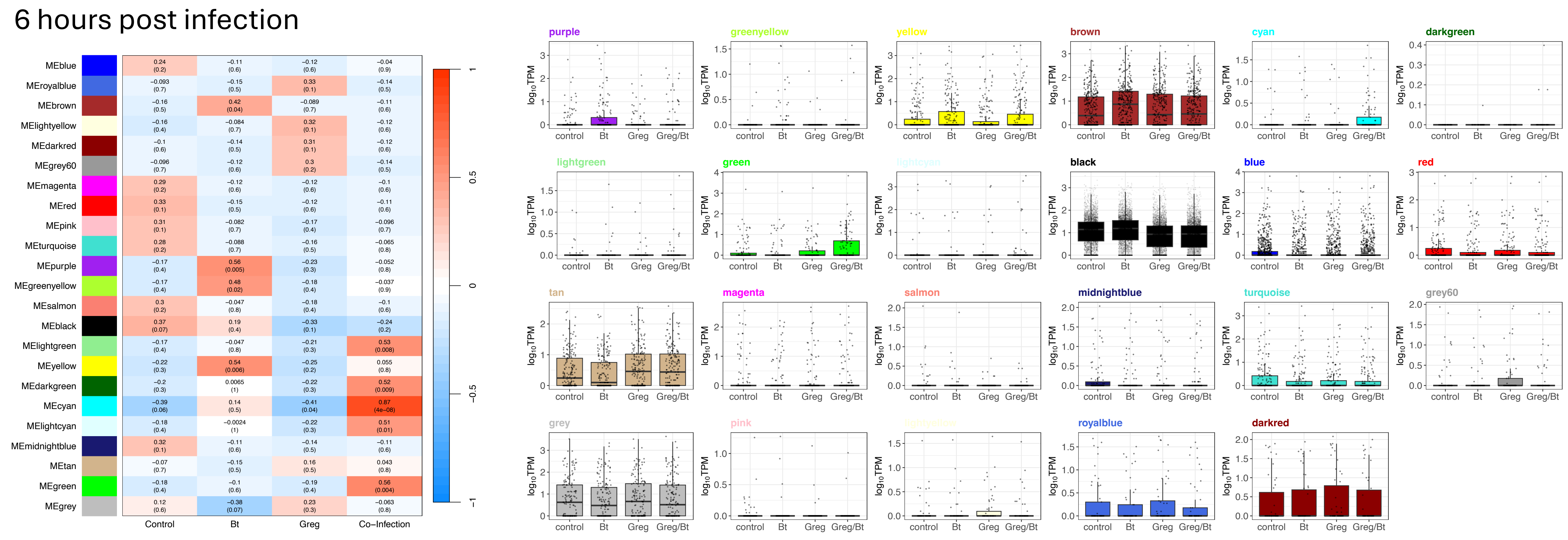
6 hours post infection
MEblue
MEroyalblue
MEbrown
MElightyellow
MEdarkred
MEgrey60
MEmagenta
MEred
MEpink
MEturquoise
MEpurple
MEgreenyellow
MEsalmon
MEblack
MElightgreen
MEyellow
MEdarkgreen
MEcyan
MElightcyan
MEmidnightblue
MEtan
MEgreen
MEgrey
Control
Bt
Greg
Co−Infection
purple
greenyellow
yellow
brown
cyan
darkgreen
lightgreen
green
lightcyan
black
blue
red
tan
magenta
salmon
midnightblue
turquoise
grey60
grey
pink
lightyellow
royalblue
darkred
log10TPM
control
Bt
Greg
Greg/Bt

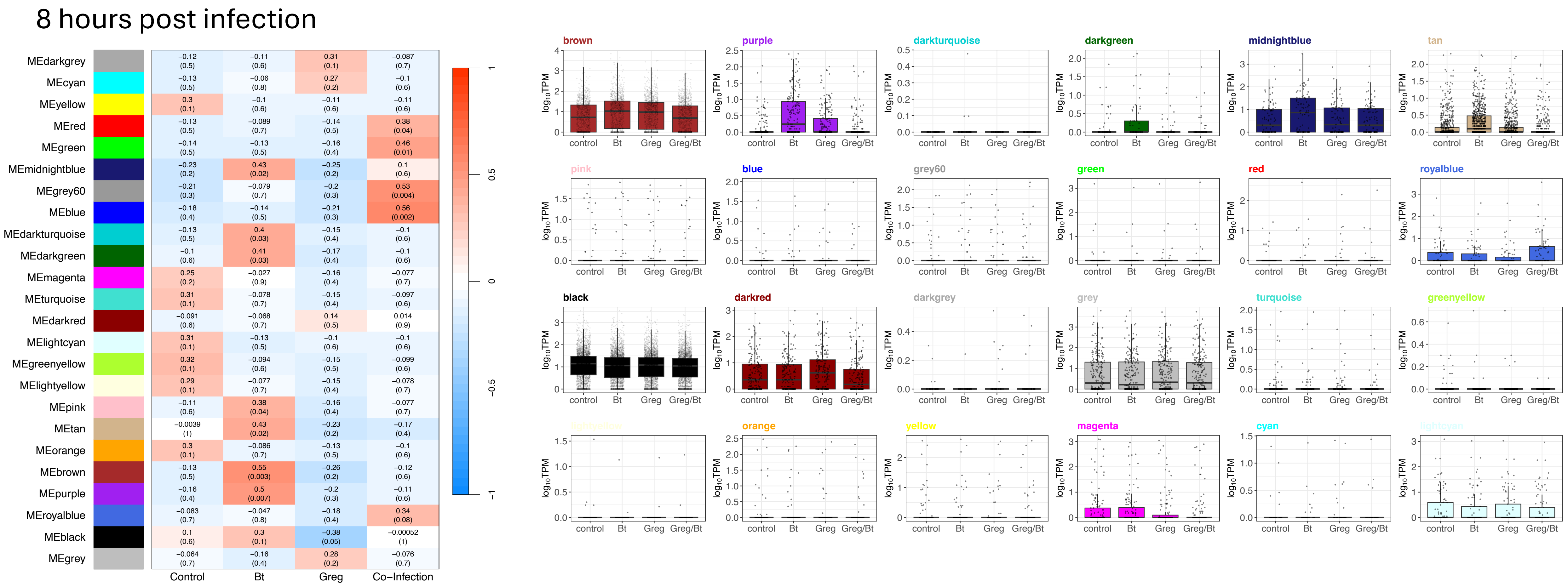
8 hours post infection
MEdarkgrey
MEcyan
MEyellow
MEred
MEgreen
MEmidnightblue
MEgrey60
MEblue
MEdarkturquoise
MEdarkgreen
MEmagenta
MEturquoise
MEdarkred
MElightcyan
MEgreenyellow
MElightyellow
MEpink
MEtan
MEorange
MEbrown
MEpurple
MEroyalblue
MEblack
MEgrey
Control
Bt
Greg
Co−Infection
brown
purple
darkturquoise
darkgreen
midnightblue
tan
pink
blue
grey60
green
red
royalblue
black
darkred
darkgrey
grey
turquoise
greenyellow
lightyellow
orange
yellow
magenta
cyan
lightcyan
log10TPM
control
Bt
Greg
Greg/Bt
